# Ancestry dynamics and trait selection in a designer cat breed

**DOI:** 10.1101/2022.12.12.520105

**Authors:** Christopher B. Kaelin, Kelly A. McGowan, Anthony D. Hutcherson, John M. Delay, Gregory S. Barsh

## Abstract

The Bengal cat breed was developed from intercrosses between the Asian leopard cat, *Prionailurus bengalensis*, and the domestic cat, *Felis silvestris catus*, with a last common ancestor 6 million years ago. Predicted to contain ~95% of their genome from domestic cats, regions of the leopard cat genome are thought to account for unique pelage traits and ornate color patterns of the Bengal breed, which are similar to those of ocelots and jaguars. We explore ancestry distribution and selection signatures in the Bengal breed using reduced representation and whole genome sequencing from 905 cats. Overall, leopard cat introgressions are reduced twofold from expectation and include examples of genetic incompatibility—reduced expression of a leopard cat gene in a domestic cat background—underlying the color traits *Charcoal* and contributing to coat color variation. Leopard cat introgressions do not show strong signatures of selection; instead, selective sweeps in Bengal cats are associated with domestic, rather than leopard cat haplotypes, and harbor candidate genes for pelage and color pattern. We identify the molecular and phenotypic basis of one selective sweep as reduced expression of the *Fgfr2* gene, which underlies *Glitter*, a desirable trait that affects hair texture and light reflectivity.

## Introduction

Interspecies hybridization as a means of introducing genetic diversity is a long-established tool for developing desirable traits in plants^1^ and animals.^2^ Most examples and studies of interspecific hybridization focus on organisms and traits that are commercially important such as tame behavior in cattle-yak hybrids^3^ or pathogen resistance in cauliflower.^4^ More recently, selection for desirable morphologic traits in companion animals has become a popular tool in the domestic cat community.

The Bengal cat breed was established in 1986 from hybrids between the domestic cat (*Felis silvestris catus*) and the Asian leopard cat (*Prionailurus bengalensis*). Leopard cats, which last shared an ancestor with domestic cats six million years ago, are similar in size to domestic cats but with a thinner body and longer hind legs (Figure 1A). The Bengal cat is now the most popular breed, with more than 200,000 registered cats. In the 35 years since breed inception, Bengal cats have been selected for the calm demeanor expected of a companion animal and morphologic traits meant to recreate or evoke the distinctive appearance of forest-dwelling felids. Such wild-like traits include a small, round head, concave nose bridge, rounded ears, and most dramatically, rosette-shaped coat color patterns characteristic of cat species like the jaguar, the margay, and the ocelot, features not found in other domestic cat breeds.

**Figure 1.**
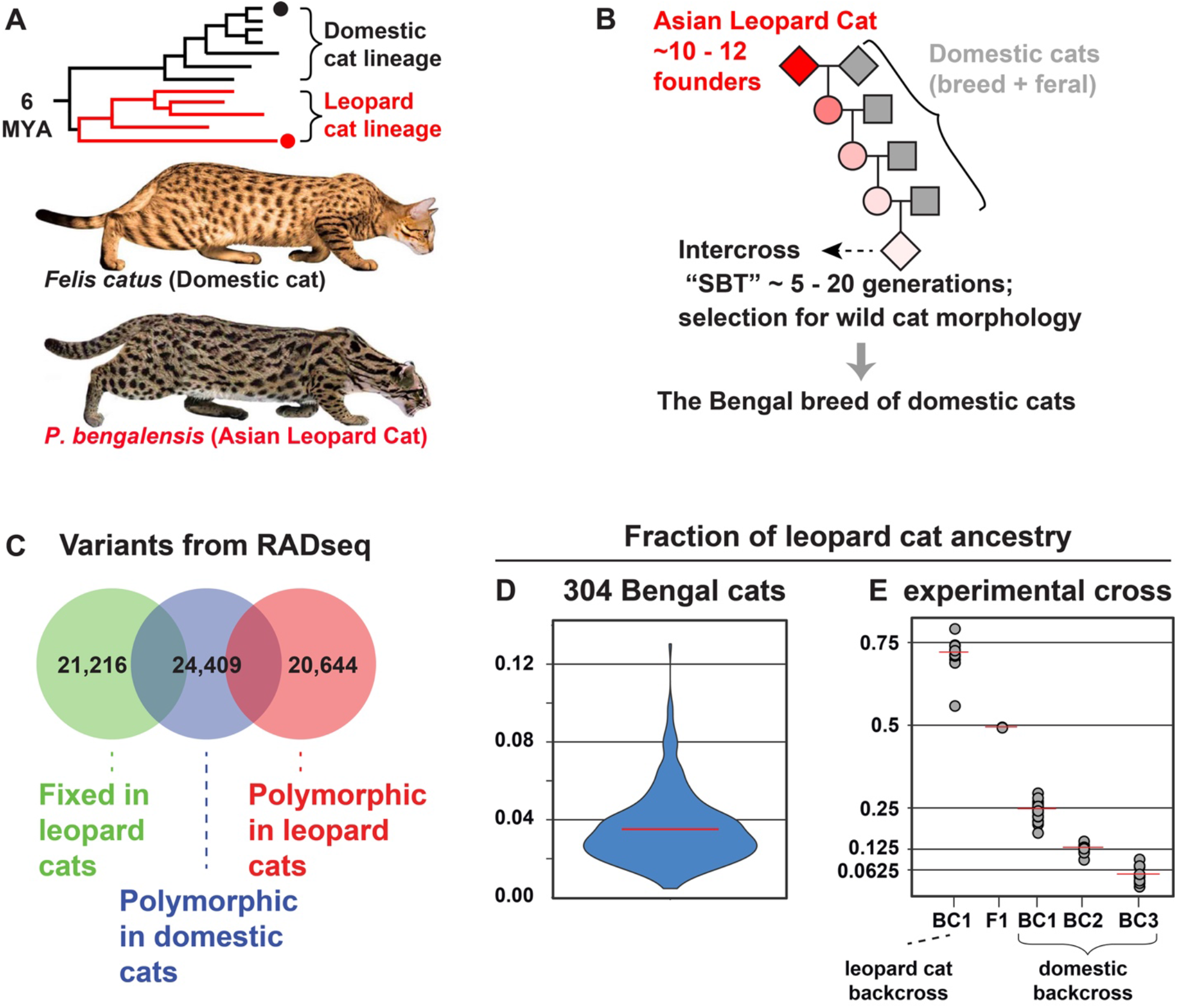
Bengal breed formation and leopard cat admixture. (A) A species tree of felid lineages that gave rise to the Asian leopard cat (*Prionailurus bengalensis*, red) the domestic cat (*Felis catus*, black). (B) A breeding paradigm for introducing leopard cat ancestry into the Bengal cat breed. (C) Distinct sets of RADseq SNVs distinguishing genetic variation within or between leopard cats and domestic cats. (D) Distribution and population mean (red bar) of leopard ancestry fraction in Bengal cats, measured from ancestry informative RADseq markers. (E) Leopard ancestry fraction in hybrids from an experimental cross, including domestic cat-leopard cat intercross offspring (F1) and subsequent backcross offspring (BC1-3). Red bars indicate mean leopard ancestry fraction per group.

With a sequence divergence of ~3% between leopard and domestic cats,^5^ the Bengal breed represents a unique community-based experiment in mammalian genetics, involving a large hybrid population characterized by disproportionate admixture between highly divergent species. As such, the Bengal breed is an excellent system with which to study genome dynamics in a hybrid population and the genetic architecture and molecular basis of their distinctive morphology, thought to be due to selection for specific regions of the leopard cat genome.

We worked with Bengal cat owners, breeders, and breed organizations to identify and quantitate regions of the leopard cat genome present in the Bengal cat breed. By giving talks at cat shows, writing articles in community newsletters, and offering to return ancestry information to breeders, we collected hundreds of buccal brush samples per year, together with bilateral photographs and pedigree information, by mail or by visiting cat shows and catteries.

Here we use restriction-associated DNA sequencing (RADseq) and/or low coverage whole genome sequencing (lcWGS) of more than 905 individuals to describe the admixture architecture in the Bengal cat breed, to identify genomic intervals under selection, and to pinpoint the genetic basis of Bengal cat traits. Within the breed, we find that leopard cat ancestry is, overall, reduced twofold from expectation and is nonrandomly distributed. We use ancestry information to identify the phenotypic consequences and genetic mechanism of one introgression—the color trait Charcoal is caused by incompatibility between the leopard cat *Agouti signaling protein (Asip*) gene and the predominantly domestic cat genome. Ancestry information also indicates that Leopard cat introgressions at *Asip* and a second gene, *Corin*, contribute to the substantial coat color diversity apparent in the breed. No leopard cat alleles, however, have swept to high frequency in Bengal cats; instead, signatures of purifying selection within the breed are associated with sequences of domestic cat origin. Genetic association reveals that a region under breed-wide selection harbors a regulatory mutation in *Fgfr2*, which arose in domestic cats and is responsible for a hair appearance and texture trait known as Glitter.

## Results

### Leopard cat ancestry in the Bengal breed

The paradigm for Bengal breed formation is to backcross F1 hybrid females to domestic cats for three generations, followed by intercrossing to similarly bred cats, which breeders refer to as stud book tradition (SBT, Figure 1B). The International Cat Association and the Cat Fanciers Association recognizes SBT cats as qualifying for breed registration. Each backcross generation dilutes leopard cat ancestry, on average, by half; thus intercrossed, breed-registered cats have an expected average of 6.25% genetic contribution from the leopard cat (Figure 1B).

We used a combination of WGS and RADseq of domestic and leopard cats to identify 66,269 single nucleotide variants (SNVs) among domestic and leopard cats within RADSeq intervals, of which 21,216 SNVs distinguish leopard from domestic cat ancestry (Figure 1C). Mean leopard cat ancestry as determined by RADSeq analysis of 304 Bengal cats was 3.48± 0.10% (range 0.46-12.3%), approximately half of that expected from the breeding paradigm (Figure 1D). We also measured leopard cat ancestry in a large intercross-backcross pedigree established in a research colony, in which the segregation of leopard cat ancestry through three successive backcross generations conformed to Mendelian expectation (Figure 1E). Thus, reduced leopard cat ancestry in the Bengal breed is not caused by transmission ratio distortion and instead is likely to represent selection against regions of the leopard cat genome and/or dilution of leopard cat ancestry by undocumented introduction of domestic cats.

Analysis of leopard vs. domestic cat ancestry by RADseq depends on genotypes at loci with a relatively uneven and sparse (1 SNV/130kbp) distribution. As an alternative approach, we used high coverage WGS for 19 leopard cats and 114 domestic cats to identify 14.6 M species informative SNVs, which allowed us to distinguish ancestry in 50kb windows by lcWGS summing allele counts across loci (Figure S1A). For 136 Bengal cats sequence by RADseq and lcWGS, estimates of genome-wide leopard cat ancestry measurements were highly correlated (Figure S1B and S1C).

The windowed approach permitted characterization of the length, number, and distribution of leopard cat haplotypes for each Bengal cat (Figure 2A) and, by extension, for the breed (Figure 2B, 2C, and 2D). Every Bengal cat we analyzed (n=737 Bengal cats) carried multiple, distinct leopard cat haplotypes, with an average of 21.4 per cat (Figure 2C). Overall, we identified 6436 haplotypes, with a mean length of 4.5Mb (range 150kb-93.3Mb, Figure 2C). There is an expectation that haplotype number increases and haplotype length decreases with successive intercross generations.

**Figure 2.**
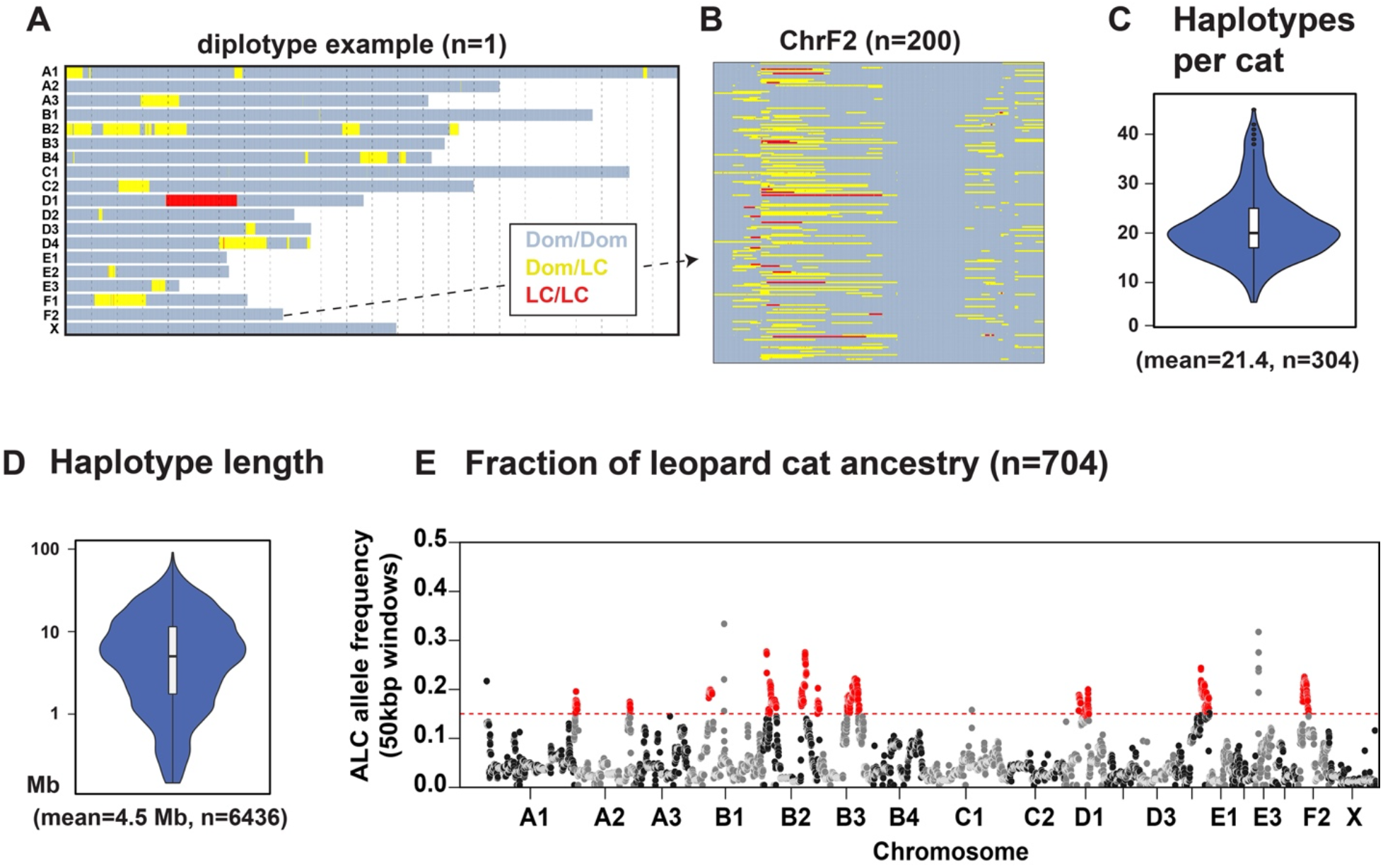
The genome distribution of leopard cat ancestry in Bengal cats. (A) A diplotype plot from a representative Bengal cat, showing species ancestry inferred from lcWGS in 48,208 50kb windows across chromosomes (in rows), colored by number of predicted leopard cat haplotypes. (B) Ancestry diplotype plot for 200 Bengal cats (in rows) across chromosome F2. (C) The distribution and quartile metrics for the number leopard cat haplotypes per Bengal cat (n=304 cats). (D) The length distribution and quartile metrics of 6436 leopard cat haplotypes inferred from 304 Bengal cats. (E) Leopard cat ancestry frequencies (n=704 SBT Bengal cats), measured from lcWGS in 48,208 50kb windows across the genome (felCat9). Windows in nine regions (red) are significantly enriched for leopard cat ancestry based on genome-wide average (Z-score > 3, red broken line).

Many leopard cat haplotypes appear to have common breakpoints, consistent with bottlenecks during breed derivation and/or selection for specific leopard cat haplotypes (Figure 2B). This interpretation is reinforced by a biased distribution of leopard cat ancestry across the genome when surveying a population of Bengal cats (Figure 2D). Quantitatively, 1,585 of 48,208 50kb windows exhibit significant deviation from a normal distribution (Z-score > 3) and appear enriched for leopard cat ancestry. Because morphologic features similar to leopard cats are prevalent in Bengal cats, we expected to find selective sweeps involving regions of leopard cat ancestry. Contrary to this expectation, the fraction of leopard cat haplotypes spanning a genomic window ranges from 0-0.27 (n=737 Bengal cats), with a median of 0.023. For windows where leopard ancestry fractions exceed 0.155, representing a Z-score > 3, nearly all (2439 out of 2448, 99.6%) cluster in 10 genomic intervals (Figure 2D).

The skewed distribution of leopard cat ancestry implies that selection and/or genetic drift had an important role in shaping ancestry architecture in the Bengal breed. But, the absence of breed-wide selective sweeps also suggests either (1) absence of strong purifying selection acting at any single locus; or (2) ongoing balancing selection. To investigate the latter possibility, we tested for excess heterozygosity, using ancestry measured in 50kb windows, from 704 SBT Bengal cats. Excess heterozygosity was detected in only 6 of 45,673 autosomal windows (chi-square test, FDR < 0.05), none of which were in the regions enriched for leopard cat ancestry, where power to detect excess heterozygosity is highest. This suggests that selection against incompatibility does not play a major role in limiting the proportion of leopard cat ancestry.

The observed architecture of leopard cat ancestry could also be a consequence of genetic drift. Low fertility and fecundity in early generation hybrids create a situation for strong genetic bottlenecks that preferentially affect leopard cat ancestry. Leopard cat haplotypes present in exceptionally fertile or desirable early generation hybrids will be overrepresented in the breed, whether or not a substrate for selection, making the role of genetic drift and selection difficult to ascertain from ancestry proportions.

The lcWGS cats represent F1 hybrid (n=7 cats), backcross (n=26 cats), and SBT (n=704 cats) generations. Excluding windows with large assembly gaps, the majority of which cluster telomeric and centromeric intervals (885 of 1007 50 kb windows, 87.9%), leopard cat ancestry is detected across 93% of the genome in SBT cats and 99.8% of the genome in the combined set of backcross and SBT cats (Figure S2A). At two regions (chrD3:17.55-36.11 Mb and chrX:35.85-40.05), leopard cat ancestry is only observed in F1 hybrids, suggesting the intervals harbor genes contributing to hybrid incompatibility, an interpretation that is supported by D3 and X chromosomes having the lowest overall proportion of leopard cat ancestry (Figure S2B and S2C).

### Genomic incompatibility underlies popular color traits

As a complementary approach to investigate how interplay between distantly related genomes has shaped the Bengal breed, we asked if specific traits observed in hybrids during breed development might have arisen from leopard cat genes. One such trait, charcoal, describes a form of partial melanism that preferentially darkens the dorsal surface, producing the appearance of a black cape along the back and mask on the face (Figure 3A). Approximately 5% of registered Bengal cats are charcoal, and a previous candidate gene study identified variants in *Agouti signaling protein (Asip*) that were associated with *Charcoal* in Bengal cats.^6^

**Figure 3.**
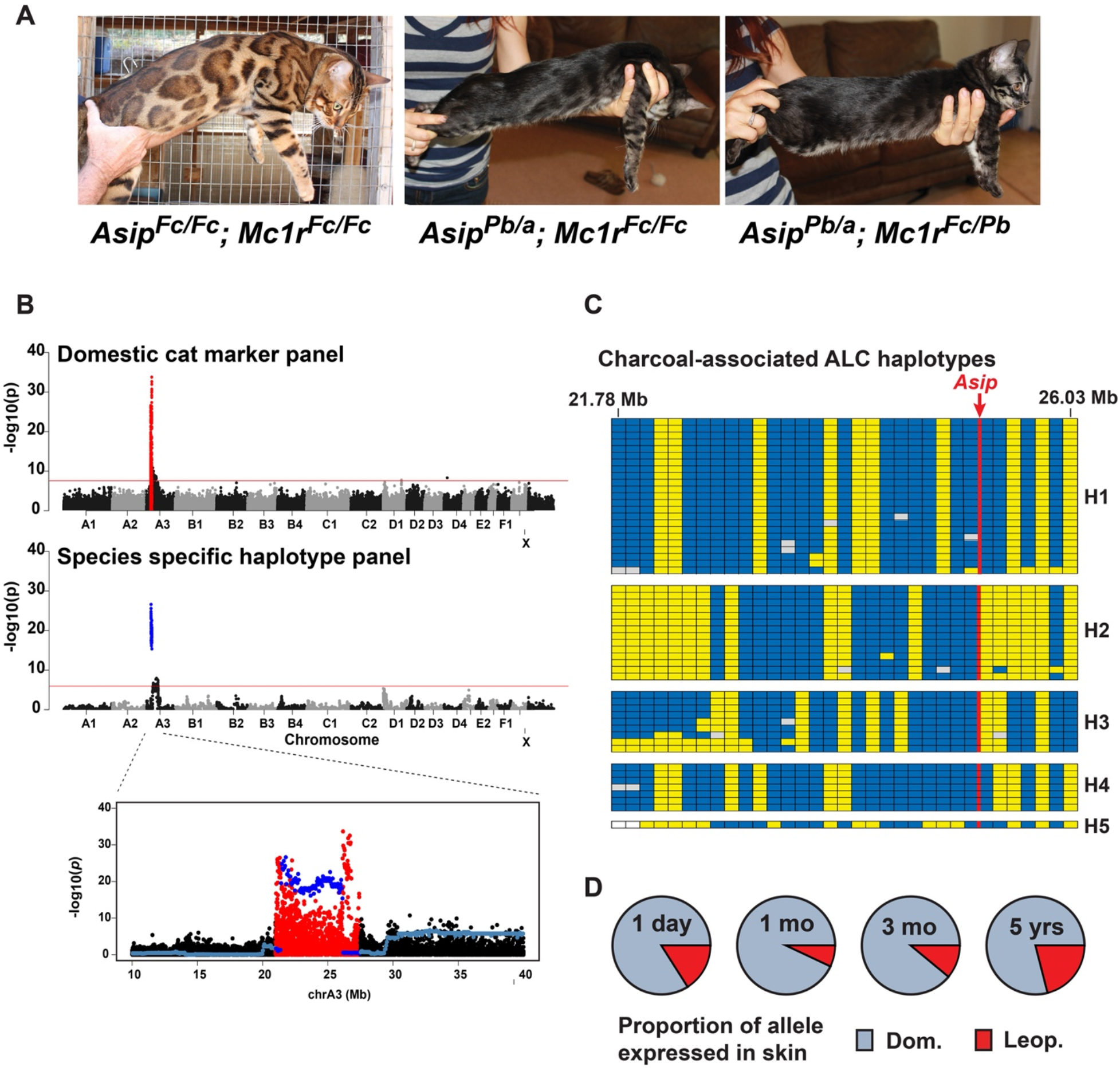
Genetic and functional characterization of *Charcoal*. (A) Images of non-charcoal (left panel) and charcoal (center and right panels) Bengal cats, with corresponding genotypes at *Asip* and *Mc1r. Pb* and *Fc* superscripts refer to the normal leopard cat and normal domestic cat alleles, respectively, and *a* refers to the *Asip nonagouti* allele. (B) Case-control GWAS for *Charcoal* using either ~1.5 million markers imputed from lcWGS (top panel) or 48,208 ancestry windows (middle panel). Significant association peaks are highlighted in red (domestic markers) or blue (ancestry windows). The relative positions of the association peaks are overlayed at higher resolution in the lower plot. (C) Five different *Charcoal*-associated leopard cat haplotypes (H1-H5), inferred from 33 leopard cat species informative RADseq SNVs. Alleles are colored as ancestral (yellow) or derivative (blue).(D) Leopard cat (*Asip^Fc^*) and domestic cat (*Asip^Pb^*) allele ratios measured by cDNA pyrosequencing from skin biopsies of four *Asip^Fc/Pb^* Bengals cats. Cat age at time of biopsy is indicated.

We carried out genome-wide association studies (GWAS) with 38 charcoal cats and 552 non-charcoal cats using ancestry inferred in 48,208 50kb windows or ~1.5M linkage disequilibrium (LD) pruned SNVs imputed from lcWGS using a domestic cat haplotype reference panel (Methods). We identified single, significant association peaks with both approaches: leopard cat ancestry in a ~5 Mb region surrounding *Asip* was significantly associated with charcoal, as were flanking domestic cat SNVs (Figure 3B), a pattern likely resulting from a recent, shared introgression event.

Five of 38 charcoal cats were homozygous for a leopard cat-derived *Asip* allele, and 27 were compound heterozygotes for a leopard cat-derived *Asip* allele and a domestic cat allele with a 2 bp coding sequence deletion, *nonagouti (Asip^a^*), that causes complete melanism when homozygous.^7^

The leopard cat *Asip* allele in charcoal cats predicts four missense alterations relative to the domestic cat protein^6^ but several lines of evidence suggested that *Charcoal* is caused by reduced expression of a normal leopard cat *Asip* gene in a domestic cat background. First, five distinct leopard cat *Asip* haplotypes, each of which can cause charcoal, are found in Bengal cats (Figure 3D). Each haplotype represents an independent introgression event from a different ancestral leopard cat chromosome. Second, a charcoal phenotype has never been reported in the leopard cat, and the missense alterations associated with the *Charcoal* allele in Bengal cats are homozygous in 21 sequenced leopard cats from three different subspecies. Moreover, the missense alterations are either ancestral in the felid lineage or represent canonical residues in other, non-melanistic mammals. Third, we identified a sibling pair of charcoal cats (*Asip^Pb/a^*), one of which also carries a leopard cat-derived allele of *Melanocortin 1 receptor (Mc1r*), which encodes the receptor for *Asip*. Both siblings have indistinguishable phenotypes (Figure 3A). Thus, a leopard cat *Mc1r* allele is unable to rescue the charcoal phenotype and indicating that reduced activity of the leopard cat *Asip* allele is unlikely to be caused by a post-translational ligand-receptor incompatibility.

We measured the levels of leopard and domestic cat *Asip* mRNA levels in 4 *Asip^Fc/Pb^* cats of varying ages. Skin punch biopsies were collected during the active phase of the hair growth cycle and allele-specific expression was determined by pyrosequencing of two polymorphic sites in *Asip* exon 2. The expression level of the *Asip^Pb^* allele was lower than the *Asip^Fc^* allele in each cat, with ratios ranging from 0.33-0.075 (Figure 3D). Reduced expression of the *Asip^Pb^* allele is likely to reflect a mismatch between domestic cat transcription factors and leopard cat cis regulatory elements. Gene regulatory incompatibilities, exemplified by the effects of *Asip* dysregulation on coat color, may contribute to selection against other leopard cat-derived genes in the Bengal cat genome.

Three additional coat color traits (colorpoint dilution, dilute, and silver) are recognized in the Bengal breed as distinct varieties, are identifiable in photographic images, and can be mapped to single genetic loci using imputed SNVs from lcWGS (Figure S3). After excluding Bengal cats with Mendelian color variants, considerable color diversity remained, ranging from grey-brown at one extreme to red-orange at the other. In 297 Bengal cats, we classified this color variation into five categories (Figure 4A) and used categorical assignment as a quantitative trait to carry out GWAS. GWAS with 48,208 ancestry windows identified broad association peaks on chrA3 and chrB1 that include all windows with *p*-value < 1×10^-3^ (Figure 4B, n=180 windows). The association intervals on chrA3 and chrB1 contain *Asip* and *Corin*, respectively, genes that act in the same genetic pathway to regulate the type (black/brown eumelanin or red/yellow pheomelanin) and amount of pigment produced in hair follicles.

**Figure 4.**
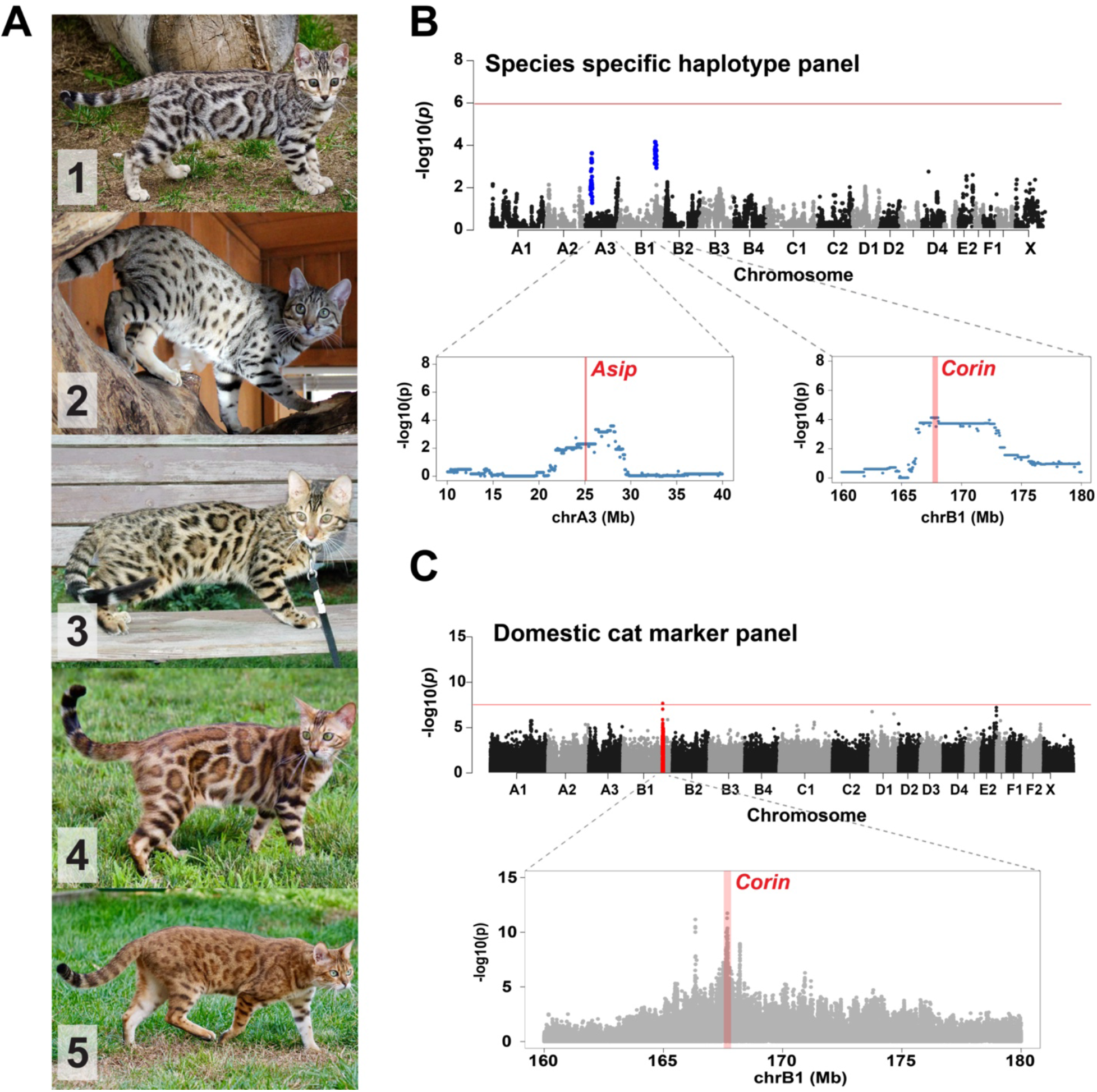
Genetic characterization of coat color diversity in Bengal cats. (A) Representative images of five coat color categories, ranging from grey-brown (category 1) to red-orange (category 5), used to classify 295 Bengal cats. (B and C) GWAS for color variation based on the scoring system in (A), using (B) 48,208 ancestry windows or (C) ~1.5 million SNV markers imputed from lcWGS. Blue (B) or red (C) association peaks are shown at higher resolution (B & C) and at higher marker density (C) in the lower panels, with the positions of candidate genes, *Asip* and *Corin*, in pink.

*Corin* encodes a Type 2 transmembrane serine protease that is co-expressed with and acts downstream of *Asip*^8^. *Corin* loss-of-function mutations, including missense mutations in domestic cats and the tiger,^9–11^ are responsible for autosomal recessive forms of golden coat color. In Bengal cats, *Asip^Pb^* and *Corin^Pb^* allele frequencies are correlated with the categorical ranking of color variation, but in opposing directions. *Asip^lc^* allele frequencies decrease and *Corin^lc^* alleles frequencies increase as coat color transitions from grey-brown to red-orange color tones across individuals (Figure 4A). Thus, leopard cat introgression is associated with loss-of-function effects at multiple loci, the combined effect of which contributes to trait diversity.

GWAS for Bengal cat color variation with ~1.5M imputed domestic cat SNVs also identified an association peak (chrB1: 167,678,371, Wald test *P* = 2.438456×10^-08^), wherein a denser marker set delineates a ~100kb association interval within *Corin* (chrB1:167,614,541-167,706,184, Figure 4C). The association interval is much narrower than the one defined by ancestry-based GWAs, and is best explained by a distinct Corin allele of domestic cat origin that also contributes to color variation in Bengal cats. None of the three *Corin* loss-of-function coding mutations responsible for recessively inherited light/amber cat color in other breeds^9,10,12^ were detected in pooled lcWGS from 737 Bengal cats. Of the 19 protein-altering *Corin* variants in Bengal cats (MAF > 3% in the pooled lcWGS, n=737 cats), 7 were among the SNVs imputed by the reference panel and were not associated with color variation. Of the remaining 12, 10 were polymorphic between leopard cats and domestic cats but not within domestic cats, and the remaining 2 were located outside the peak interval of association, suggesting that a domestic cat SNV affecting color diversity is likely regulatory.

### Domestic cat ancestry in the Bengal breed

For most domestic cats (and dogs), morphologic diversity between breeds far exceeds diversity within breeds. The Bengal cat breed is an exception because many of the domestic cats used in Bengal breed foundation were not from registered breeds, but were instead feral cats, providing an opportunity for positive selection of domestic cat-derived alleles that might be unique to the Bengal breed.

We searched for regions that might represent positive selection for domestic cat alleles in Bengal cats by examining the intersection between regions of low within-breed diversity and regions of high between-breed diversity. Starting with a panel of 19.6 million SNVs identified from high coverage WGS in 26 domestic cats, we identified regions with low within-breed diversity in lcWGS from 387 Bengal cats by calculating pooled heterozygosity (H_p_) in discrete windows across the genome. We tested different window sizes to minimize H_p_ variance and maximize resolution, settling on 11,468 200 kb intervals covering all assembled chromosomes. 180 intervals (1.57%) had a Z-score < −3, indicating significant enrichment for low diversity regions based on Gaussian expectation. 81.6% (147/180) of these low diversity windows cluster in groups of 2 or more windows within 21 genomic regions, ranging in size from 0.4-10 Mb (Figure 5A).

**Figure 5.**
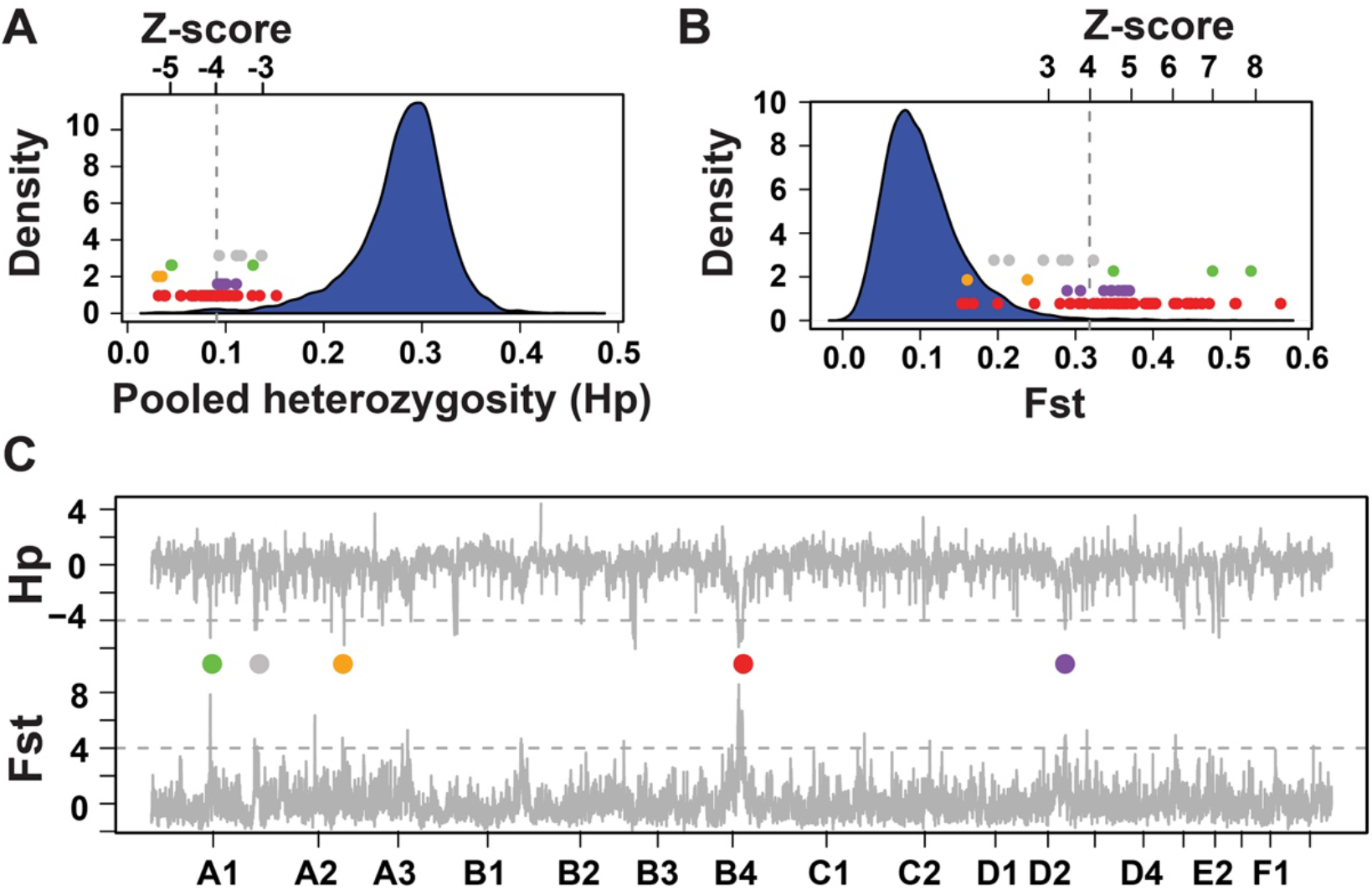
Evidence for selection in the Bengal cat genome. The frequency distributions of (A) pooled heterozygosity (H_p_) from 387 Bengal cats sequenced by lcWGS and (B) the fixation index (FST) between Bengal cats and other domestic cats, measured in (C) 11,468 200 kb windows across the genome. The colored circles indicate cluster of windows with both low H_p_ (Z-score < −3) and high FST (Z-score > 3).

To identify regions of high between-breed diversity, we calculated the fixation index (Fst) in 200kb intervals between the Bengal population and the domestic cats used for variant discovery. Of the 11,468 200 kb windows, 160 (1.40%) had an FST Z-score > 3, and 117 windows (73.1%) (117/160) were clustered in groups of two or more within 17 genomic intervals, ranging in size from 0.4-10.8 Mb (Figure 5B). 73 windows exhibited both low H_p_ (Z-score < −3) and high FST (Z-score > 3), of which 63 were clustered in five genomic intervals, ranging in size from 0.4-10 Mb (Figure 5C).

### Selection and the genetic and functional basis for *Glitter*

In considering the potential phenotypic underpinnings of the candidate positive selection regions depicted in Figure 5C, our attention turned to glitter, a coat condition that softens hair texture and causes an iridescent sheen to the coat (Figure 6A). Breeders consider glitter to be a desirable, breed specific trait with an autosomal recessive mode of inheritance.

**Figure 6.**
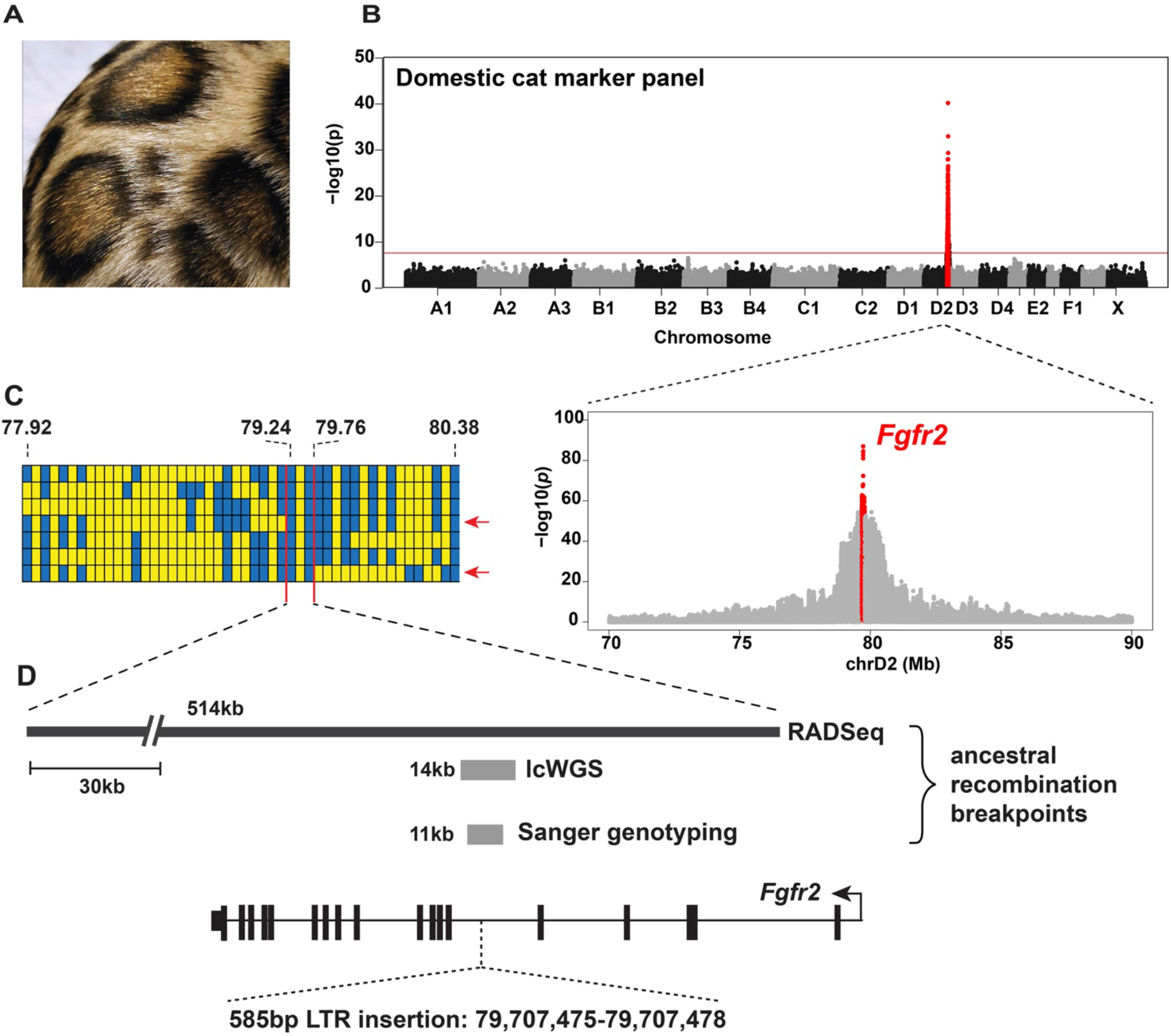
Genetic characterization of *Glitter* in Bengal cats. (A) The iridescent sheen apparent in the coat of glitter cats. (B) Top panel - a case-control GWAS to identify a *Glitter* association interval (red), using ~1.5 million lcWGS-imputed SNVs, with an association peak at chrD2:79,620,465 (Wald test *P* = 9.0×10^-41^). Bottom panel – a higher resolution view of the *Glitter* association interval with increased marker density and SNVs located within the candidate gene, *Fgfr2*, highlighted in red. (C) Recombinant *Glitter* haplotypes (in rows) inferred within a 2.57Mb interval, defined by association with RADseq markers (in columns, Figure S4), and colored by reference (yellow) or alternate (blue) allele in felCat9. (D) *Glitter* genomic interval refinement, using inferred ancestry recombinant breakpoints from both RADseq and lcWGS (see methods) to identify 14,229 bp and 10,493 bp genomic intervals (grey bars) with a Bengal cat specific LTR insertion in *Fgfr2* intron 5 at chrD2:79,707,475-79,707,478.

Independent of selection scans, we carried out a case-control GWAS for *Glitter*. From photographs submitted with DNA samples, we could confidently distinguish 371 glitter and 188 non-glitter cats from lcWGS genotyped cats, and 144 glitter and 82 non-glitter individuals from RADseq genotyped cats. Both approaches identified a broad, 2.47 Mb association interval (chrD2:77,904,283-80,383,333), that comprises 13 genes and overlaps one of regions under purifying selection in the Bengal cat breed (Figure 6B and S5A). Evaluating all imputation loci revealed a highly significant association peak within intron 5 of *Fibroblast growth receptor 2 (Fgfr2*) (chrD2:79,714,539 and 79,715,317, Wald test *P* = 1.03×10^-87^, Wald test), which we considered a strong candidate based on prior studies.^13^

We inferred haplotypes across the *Glitter* association interval using 48 loci corresponding to domestic cat polymorphic markers. By contrast to the situation with *Charcoal*, all 144 glitter cats shared a single haplotype of 513,890 bp (chrD2:79244423-79758313) (Figure 6C). This interval overlaps a single gene, *Fgfr2*.

To identify the *Glitter* mutation, we used lcWGS to extend haplotype mapping (Methods) followed by direct genotyping of individual cats to delineate a core haplotype of 10,985 bp shared among 5 recombinant and 1 non-recombinant glitter cats (Figure 6C). There are 18 SNVs and 5 indels in this region, but none of them were strong candidates for causal variants based on allele frequency in other domestic cats and inferred deleteriousness.

However, in lcWGS data from *Glitter* cats we identified a 585 bp ERV1-3_FCa-type long terminal repeat (LTR) insertion within the *Glitter* interval (Figure 6D). Among 57 domestic cats for whom high coverage WGS was available, the LTR insertion was only present in Bengal cats, and two additional breeds, Egyptian Maus and Toyger’s, in which glitter was introduced through shared ancestry with Bengal cats. We genotyped the LTR insertion with a PCR-based assay in 133 Bengal cats, including 86 glitter and 47 non-glitter cats. All 86 glitter cats were homozygous for the LTR insertion, while all non-glitter cats were either heterozygous or homozygous for the reference allele (Table S1). As described below, there is strong evidence that *Glitter* is a hypomorphic mutation of *Fgfr2* and in what follows we refer to the two alleles as *Fgfr2^gl^* and *Fgfr2^+^*.

Histologically, we observed two abnormalities in hairs from *Glitter* animals (Figure 7A and 7B). For all four types of hair (guard, awn, down, and awn-down), the width of the medulla in *Fgfr2^gl/gl^* hair is significantly reduced compared to *Fgfr2^gl+^* or *Fgfr2^+/+^* hair (Figure 6A). Reduced medullary width is also associated with a similar trait called *Satin* in mice, caused by loss-of-function for *Foxq1*,^14^ and in rats, for which *Fgf7* was suggested as a candidate gene.^15^ Reduced medullary width may explain differences in hair texture but is unlikely to account for the characteristic appearance for which both *Glitter* and *Satin* are named. We noticed, however, an additional abnormality in down hairs, which account for 75% of the coat.^16^ Normally, the hair medulla contains a vertical array of keratinocytes that alternate with vacuolated air cells,^17^ and in down hairs this array becomes thin and fragmented towards the tip of the hair (Figure 7B). In *Fgfr2^gl/+^* or *Fgfr2^+/+^* hair, the gap between the thin, fragmented portion of the medulla begins ~1mM from the distal tip but in *Fgfr2^gl/gl^* hairs, the gap is much larger (mean 3.8mM, Student’s t-test *P* = 1.9×10^-14^), and likely explains the translucent and iridescent appearance of the glitter coat.

**Figure 7.**
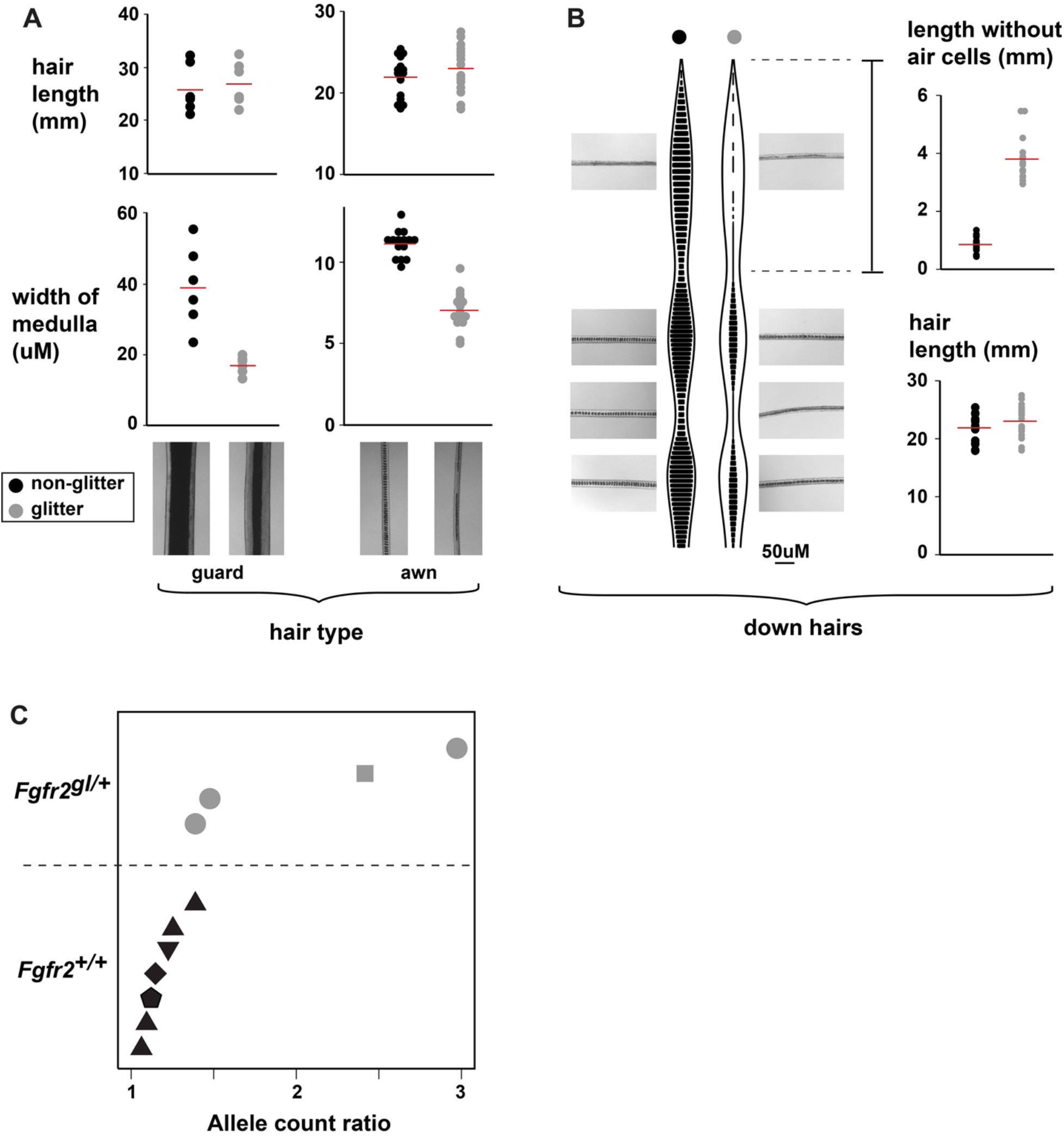
The functional basis of glitter in Bengal cats. (A) Measures of hair length and medulla width, in guard (left panels, n=6 hairs per group) and awn hairs (right panels, n=15 hairs per group), from three non-glitter (black) and three glitter (grey) Bengal cats, with mean values per group (red lines). Representative images for each hair type are shown. (B) Cartoon representation and photographic images of at different positions along the length of down hairs from non-glitter (left, black circle) and glitter (right, grey circle) Bengal cats. Measures of hair tip medulla fragmentation (top right) and hair length (n=15 hairs from 3 Bengal cats in each group) in down hairs. (C) Allele count ratios at *Fgfr2* polymorphic sites in transcripts from whole skin RNAseq. Symbols represent different cats, 2 *Fgfr2^gl/+^* (grey) and 3 *Fgfr2^+/+^* (black). Ratios in *Fgfr2^+/+^* samples are included to reflect a normal range of *Fgfr2* allelic variation.

Increased expression of *Fgfr2* is proposed to contribute to increased hair thickness in humans,^18^ and a dominant negative *Fgfr2* transgene causes reduced medullary width in transgenic mice.^13^ We measured the impact of the LTR insertion on *Fgfr2* expression in skin biopsies from two *Fgfr2^gl/+^* Bengal cats, in which allele-specific expression could be monitored by counting RNA-seq reads that overlapped 4 polymorphic variants. The ratio of *Fgfr2^+^* to *Fgfr2^gl^* varied from 1.4 – 3.0 (mean, 2.1) across the 4 variants (Figure 7C). By contrast, the “normal” range of allelic ratios in 3 *Fgfr2^+/+^* samples (using 7 polymorphic variants) was 1.1 – 1.4 (mean, 1.2). We conclude that *Glitter* is caused by reduced expression of *Fgfr2* that leads to changes in hair thickness and structure without apparent effects on other aspects of morphology or fitness.

Notably, a complete loss-of-function for *Fgfr2* is embryonic lethal in mice,^19^ while gain-of-function mutations are well-recognized causes of severe Mendelian craniofacial conditions.^20^ More comprehensive analyses will be needed to rigorously determine the impact of the *Glitter* LTR insertion on transcript isoform patterns and amounts, but it seems likely that *Glitter* homozygotes have more than a 50% reduction in *Fgfr2* activity since hair defects have not been described in mice that are heterozygous for gene-targeted loss-of-function alleles.^19^ In this regard, *Glitter* is somewhat analogous to naturally occurring mutations of other pleiotropic regulators that are targets of natural and/or artificial selection.

### Concluding remarks

The potential for hybridization and exchange of discrete genomic segments between species is being increasingly recognized as an important mechanism for adaptation in a wide range of species, including butterflies,^21,22^ Darwin’s finches,^23^ fish,^24,25^ humans,^26,27^ and felids.^28^ Such processes can be difficult to study and model because of limited access to samples and/or the challenges of modelling demographic history. Bengal cats provide some unique advantages to study inter-species introgressions: (1) initial hybridization is documented and breed history is relatively short and controlled; (2) the hybridized species are genetically distinct, sharing a common ancestor more than six million years ago; (3) there is extensive phenotypic diversity among a large readily accessible population; and (4) morphologic features of leopard cats are represented and selected for in Bengals providing a number of opportunities to understand how and to what extent introgression events contribute to phenotypic variation.

Bengal cats are also unique in that they represent a community-based effort driven by popularity of and interest in companion animals, taking advantage of a highly branched felid phylogeny in which chromosome number and structure has been largely conserved. Development of new breeds of domestic cats based on distantly related wild x domestic cat hybrids is being actively pursued by many cat fanciers and will likely prove useful to scientists as well as cat owners.

## Acknowledgments

The authors thank Hermogenes Manuel for technical assistance, Karen Sausman, Jon and Robyn Patterson, Teresa Seling, and many other members of The International Cat Association for assistance with sample collection. This research has been supported in part by the HudsonAlpha Institute for Biotechnology.

## Authors contributions

C.B.K., K.A.M., and G.S.B. conceived the project, designed experiments, and wrote the paper; A.D.H. and C.B.K. collected samples and analyzed data. J.M.D analyzed data; G.S.B. procured funding.

## Methods

### Sample collection and processing

Buccal swab samples were collected at cat shows and catteries, or by mail submission directly from breeders, using either CytoPak brushes (Medical Packaging Corporation) or cotton-tipped wood applicators (Dynarex). For whole blood collection, samples were obtained by veterinary staff in EDTA-coated (purple cap) tubes. Tissue samples, from spay/neuter surgeries or after euthanasia, were collected in conical tubes and preserved in RNAlater (Thermo Fisher). DNA extractions from buccal swabs and blood was carried out using the QiaAmp DNA extraction kit (Qiagen), following the manufacturer’s instructions, and eluted in 100ul. For tissue samples, ~50mg tissue samples were extracted in lysis buffer with Proteinase K, followed by phenol-chloroform extraction, ethanol precipitation, and resuspension in 200ul of Tris-buffered EDTA. All DNA samples were stored at −80C.

Dorsolateral whole skin isolates for RNA extraction were obtained using 4mm biopsy punches (Acuderm Inc.), after application of local anesthetic. Hair surrounding the biopsy site was shaved seven days prior to biopsy collection to reset the hair growth cycle. Paired biopsies within and outside plucked areas were visually inspected under a dissecting scope to verify that biopsies from shaved regions were in active hair growth. Two skin samples were also obtained from neonatal kittens euthanized for unrelated health issues. Skin biopsies (10-20mg) were homogenized in 2mLs of Trizol using a Polytron rotor-stator followed by RNA isolation using TRIzol Plus RNA Purification Kit (Thermo Fisher).

We constructed SbfI-restriction site assisted reduced representation DNA sequencing (RADseq) libraries for 578 cats, following a protocol by Etter et al., ^29^ and sequenced them on either GAIIx or HiSeq Illumina platforms. The sample set includes 369 Bengals cats, 71 cats from an experimental domestic cat-leopard cat intercross pedigree (Figure 1E), 47 cats from multiple breeds that contributed to Bengal breed development, 80 feral cats, and 11 leopard cats. Two additional Asian leopard cats sequenced to >20x coverage were also included to create a species informative marker panel (see below).

### RADseq marker panels curation and genotyping

We established three non-overlapping sets of markers to analyze genetic variation and admixture in the Bengal cat breed (Figure 1C): (1) SNVs that are polymorphic in domestic cats but not leopard cats, (2) SNVs that are polymorphic in Asian leopard cats but not domestic cats, (3) SNVs fixed for different alleles in domestic cats and leopard cats.

To identify and classify these markers, we aligned sequencing data, performed variant calling, and compared variant sets from leopard cat and domestic cat genomic data. We aligned leopard cat RADseq with BWA to the domestic cat assembly (felCat8) and performed joint variant calling with Freebayes to identify 92,356 biallelic SNVs, 71,943 of which had a non-reference allele frequency > 0.08 and 50,662 of those were represented in at least 70% (8 of 11) of individuals. We also trimmed and aligned WGS from 26 domestic cats (15 random bred cats and 11 breed cats), to felcat8 with Cutadapt and BWA-mem, respectively. We then used Platypus and VCFtools to identify and filter 19,589,252 biallelic SNPs, resulting in 31,55 SNVs within RADseq intervals.

The set of 50,662 and 31,554 variants form leopard and domestic cats, respectively, were reciprocally filtered with BEDtools to filter overlapping loci. SNVs in leopard cats were further distinguished as fixed (non-reference allele frequency > 0.93) or polymorphic (non-reference allele frequency < 0.93). We used UCSC LiftOver tool to convert genome coordinates from felCat8 to felCat9, retaining 99.5% of loci. The filtering and thresholding steps result in 66,289 SNVs that could be split into three non-intersecting marker sets capable of distinguishing species of origin and inferring domestic and leopard cat haplotypes, including: 21,216 SNVs fixed for different alleles between species, 20,664 SNVs polymorphic only in leopard cats, and 24,409 SNVs that are polymorphic only in domestic cats.

RADseq from 578 cats was aligned to the domestic cat assembly (felCat9) with BWA, and genotypes were called at the 66,289 marker loci with Freebayes.

### Ancestry inference in domestic cat hybrids

We inferred species ancestry in hybrid cats with >10x mean coverage depth, including 300 Bengal cats and 72 cats from the hybrid backcross pedigree. To estimate the fraction of leopard cat ancestry per cat using RADseq genotypes, we summed leopard cat alleles at all autosomal loci and divided by the total number of autosomal alleles (2x the number of genotyped loci).

To detect, visualize, and characterize leopard cat haplotypes in the Bengal cat genome, we used BEAGLE^30^ to impute missing genotypes and VCFtools to generate a genotype matrix, with genotypes coded as 0, 1, or 2 based on the non-reference (leopard cat) allele count. The matrix was used to visualize leopard cat introgression haplotypes in diplotype plots (Figure 2A) using a custom R script and to characterize haplotype number (Figure 2C) and length (Figure 2D) using a custom python script.

### GWAS in Bengal cats using RADseq genotypes

We used GEMMA (v0.98.3)^31^ to perform GWAS with RADseq genotype data converted from VCF to Plink BED format, after filtering cats with mean read depth < 6. We calculated a centered relatedness matrix with GEMMA’s built-in function using 24,409 domestic cat specific SNVs. GWAS were then performed using the estimated relatedness matrix on all 66,289 RADseq SNVs, using GEMMA linear mixed model function and a Wald test for association. GWAS were plotted with the qqman package in R.

### A genome-wide marker panel for ancestry inference from lcWGS

To develop a dense marker panel for ancestry inference, we trimmed and aligned ~10x WGS from 19 leopard cats (PRJNA348661) to the domestic cat genome (felCat9). After marking duplicate reads with Picard, joint variant calling with the Sentieon pipeline identified 78,311,729 variants. Applying standard filters, we identified 19,361,272 SNVs that were fixed or nearly fixed (non-reference AF > 0.95) in the leopard cat, and 12,016,231 SNVs that were polymorphic (MAF > 0.1) among the genotyped leopard cats.

Domestic cat variants were filtered from a 99Lives VCF file with 284 cats called against felCat9^32^. We first filtered to exclude individuals likely to have non-domestic cat ancestry and retained 114 cats with 35,919,989 variants (MAF > 0.01). To establish a set of SNVs that are fixed for different alleles in the domestic and leopard cat, we filtered the 19,361,272 leopard cat SNVs to remove sites that overlap with the domestic cat variants, resulting in 14,631,687 SNVs that predict species ancestry.

### Low coverage whole genome sequencing

We constructed lcWGS libraries used a modified version of a protocol developed by Baym et al. (2015),^33^ in which Nextera DNA library prep reagents are diluted to reduce library preparation cost. Briefly, we constructed indexed library pools, in 96-well format, from 2.5 ng of genomic DNA, using a 10-fold dilution of Nextera DNA Library Prep Kit reagents. Libraries were pooled by row (n=12) after initial amplification and pooled by plate before Illumina sequencing. 838 Bengal cat libraries were sequenced to generate 150PE reads on HiSeq or Novaseq platforms to an average depth of 4.48 million read pairs per library.

Demultiplexed lcWGS was trimmed with CutAdapt and aligned to felCat9 with BWA-mem. Overlap between paired end reads was removed using the clipOverlap function (BAM tools), and duplicate reads were marked using Picard. Alleles at 14,631,687 species informative SNVs were counted using the Platypus and VCFtools. Allele counts for reference (domestic cat) and alternate (leopard cat alleles) were summed in 50kb non-overlapping windows using BEDtools. For each window, ancestry was determined as the ratio of domestic cat and leopard cat allele counts.

To identify empiric thresholds for ancestry inference, we used the above workflow to assign ancestry in 50kb windows to four Bengal cats sequenced at >20x mean coverage depth. At 20x depth, a trimodal distribution of allele ratios, centered at 0, 0.5, and 1, is observed for 48,208 50kb windows (Figure S1A). The modes correspond to windows that are homozygous for domestic cat ancestry, heterozygous, and homozygous for leopard cat ancestry, respectively.

We then downsampled aligned reads to mean read depths ranging from 1x to 0.05x. Trimodal distributions could be delineated down to 0.1x coverage depth, with inflection points between modes occurring at 0.15 and 0.80 (Figure S1A). We also tested different window sizes, ranging from 10kb to 200kb (data not shown), and 50kb windows provided an optimal balance, allowing for accurate ancestry inference for coverage depths achieved with lcWGS (between 0.1-1x) and sufficient resolution to detect leopard cat admixture given the distribution of leopard cat haplotype length in Bengal cats. This approach was used to infer ancestry in 48,208 50kb windows across the genome in 737 Bengal cats with lcWGS.

### ALC ancestry measures

The percentage of ALC ancestry per individual Bengal cats, was measured from lcWGS in the following way. Genomic intervals were filtered to include only 50kb windows in assembled autosomes (46,603 windows). Additional filtering excluded 990 windows (2.0% of assembled autosomes), without species information markers. These windows overlap large assembly gaps, predominantly at centromeres, in the felCat9 assembly (Figure S2A). The remaining 45,613 windows were used to estimate the amount ALC ancestry for each Bengal cat, by summing species informative allele counts to infer haplotype counts within each 50kb window, and then determining the genome wide ALC ancestry as total ALC haplotype counts divided by 2 * the number of ancestry windows.

To assess the precision of leopard cat ancestry estimates by RADseq and lcWGS, we independently measured ancestry for 136 Bengal cats genotyped using both methods. Leopard cat ancestry fractions from the same individual were correlated (Pearson *r* = 0.9183) and mean ancestry did not differ significantly (lcWGS: 3.43±0.15%, RADseq: 3.33±0.15%; Student’s *t*-test *P* = 0.104, Fig.S2) between RADseq and lcWGS estimates.

We also estimated breed wide ALC ancestry in 50kb windows across the genome, using the 737 Bengal cats (Fig. S3A). The cohort of Bengals cats includes seven F1 interspecies hybrids, eight N2 backcross offspring, and 722 cats with ALC ancestry from 0.4% to 16%, which predominantly represent the interbreeding population of SBT Bengal cats, as well a few N3 backcross offspring from foundation breeding.

The assembled portion of felCat9 spans 49,215 50kb windows, 1007 of which contain no (or relatively few) species informative markers because they overlap large assembly gaps, predominantly at centromeres. This leaves 48,208 windows within which to assess ALC haplotype count ratios from all 737 Bengal cats. This was accomplished by first determining ALC haplotypes counts per window in each cat, and then using population data to summarize the fraction of ALC haplotype counts per window across the populations, for each window across the genome (Fig. S3).

### Genotype imputation from lcWGS

To impute genotypes from lcWGS, we used an imputation panel and variant calling pipeline developed by Gencove for the domestic cat. Fastqs from 737 Bengal cat samples and 69 cats from 5 additional breeds were uploaded to Gencove, Inc. using the Gencove command line interface. Imputation was performed using loimpute as part of the Gencove imputation pipeline, using the domestic cat v2.0 reference panel with 187 domestic cats. VCF files for each cat with genotype calls at 55,210,041 sites were subsequently merged, annotated, and filtered to include only SNVs at MAF > 0.05 in 704 SBT Bengal cats, using Bcftools.

For downstream analyses, VCF files were converted to binary Plink format and pruned using Plink’s variance inflation factor method (SNP window = 50, SNP window step = 5, variance inflation factor = 2) to extract 1,583,065 LD-pruned SNVs.

To determine imputation accuracy, we measured concordance rates at 23,877 SNVs genotyped by both RADseq and lcWGS imputation in 40 Bengal cats. RADseq genotypes, considered a “truth set”, were filtered to include (1) cats with >20x mean coverage depth, (2) sites with >20x coverage depth, resulting in between 10,402 and 21,346 sites evaluated per cat. After filtering, genotype concordance, stratified by RADseq genotype classification (homozygous reference, heterozygous, and homozygous alternate), was measured as the number of lcWGS imputation genotypes that match RADseq genotypes (Fig. SX).

### GWAS with lcWGS imputation

Imputed genotypes from low pass sequencing were used to characterize breed structure within Bengal cats and in relationship to 5 additional breeds – Abyssinian, Egyptian Mau, Ocicat, Singapura, and Toyger, using Plinks PCA function and plotting in R.

GWAs were carried out using the linear mixed model approach implemented in Gemma (v 0.98.3). Pruned variants (Plink1.9) were used to estimate a relatedness matrix and as input for a univariate linear mixed model with Wald test. GWAs were plotted in R using qqman package.

### *Glitter* interval refinement

GWAs for *Glitter* were performed independently using either imputed lcWGS (Figure 5) or RADseq markers (Figure S4). For RADseq data, we could confidently evaluate the glitter trait in 226 of 323 images, including 144 glitter and 82 non-glitter individuals. We performed a case-control GWAs with Plink using 66,269 SNVs (Figure 1C), assuming a recessive model. Genome-wide and locus specific association was plotted using qqman package in R.

We next used Beagle to impute missing RADseq genotypes and infer haplotypes across chromosome D2 using domestic cat markers (1368 RADseq SNVs), in 226 Bengal cats and 30 cats from other breeds. Among 146 glitter Bengal cats, 221 haplotypes were identical across the 2.47 Mb *Glitter* association interval (48 RADseq SNVs). The remaining 71 haplotypes were presumed to represent ancestral recombination breakpoints, of which 2 on each side define a 513,891 bp (chrD2:79244423-79758313, felCat9) *Glitter* genetic interval.

We then used lcWGS to refine the *Glitter* interval and identify candidate causal variants. We could determine glitter status for 200 of 287 lcWGS-sequenced Bengal cats, of which 131 were glitter and 69 were non-glitter. lcWGS was pooled according to *Glitter* status to permit variant discovery and allele frequency estimates for each pool. Using Freebayes (v1.3.1, Freebayes -m 15 -q 15 -C 1 -G 2 --maxcoverage 250), we identified 3408 and 6725 polymorphic sites within glitter and non-glitter pools, respectively. Because the *Glitter* haplotype is expected to be present in both pools, we focused on 2936 shared variants within *Glitter* interval. At these sites, minor allele frequencies were significantly lower in the glitter pool, and underrepresented alleles were associated with only 12 of 131 glitter individuals. We interpreted these observations as evidence of recombinant haplotypes with breakpoints within the current interval that delineates14,240 bp interval within the *Fgfr2* transcriptional unit. Sanger sequencing of 13 amplicons in 8 Bengal cats (2 non-glitter and 6 glitter) confirmed haplotype breakpoints on either side of the *Glitter* interval and further refined the interval to 10,494 bp.

### Measurement of allelic expression

For *Asip* allele ratio measurements, cDNA was prepared from 500ng of total RNA using Superscript IV Reverse Transcriptase (Thermo Fisher). Pyrosequencing was performed using a PyroMark Q24 DNA Sequencer (Qiagen). PCR primers (Forward: 5’ – GAGCAACTCCTCCATGAACAT – 3’ and Reverse: 5’ – CGAAGCCTTTTTCTTGGAAGATC – 3’) and sequencing primer (5’ – TTTGGATTTCTTGTTCAG – 3’) target cDNA region with two species informative SNVs (c.T142C and c.G162A) used for allele ratio measures.

(Note: in addition to collecting biopsies in areas with plucked hairs, we also collected biopsies in adjacent areas. These biopsies were included in the analysis. In all cases, Asip allelic expression ratios are closer to 1, suggesting that allele biased expression is more pronounced during active hair growth, when Agouti expression is maximal.)

For *Fgfr2* allele ratio measurements, strand-specific RNA sequencing libraries were prepared from two neonatal dorsal skin biopsies, each heterozygous for the Glitter-associated LTR insertion (*Fgfr2^g/+^*, Table S1). The libraries were sequenced to 100M reads (50SE, Illumina HiSeq2500) and 33M read pairs (150PE, Illumina Novaseq6000) for Bgl614 and Bgl5781, respectively. Reads were aligned to the domestic cat genome assembly (Felis_catus_9.0) with NCBI *Felis catus* annotation release 104 using the STAR aligner and read counts overlapping SNVs were used to infer allelic transcript ratios. Three and one SNV discriminated allelic transcripts for Bgl612 and Bgl5781, respectively.

### Hair morphometry

Hairs from three non-glitter and three glitter Bengal cats were plucked by cat breeders. Hairs were mounted in mineral oil between a glass slide and coverslip. Photomicrographs were tiled along the length of five down hairs and two guard hairs from each animal. Hair length, area of the hair and area of the medulla was measured in each tiled image using ImageJ (version 1.45s).^34^ The average hair width and average medullary width were computed: average hair width = total area of the hair/total hair length and average medullary width = total area of the medulla/total hair length. The length of the hair without air cells was measured from the tip of the hair on down hairs.

