## Supplemental Information for "Ancestry dynamics and trait selection in a designer cat breed"

**Christopher B. Kaelin, et al.**

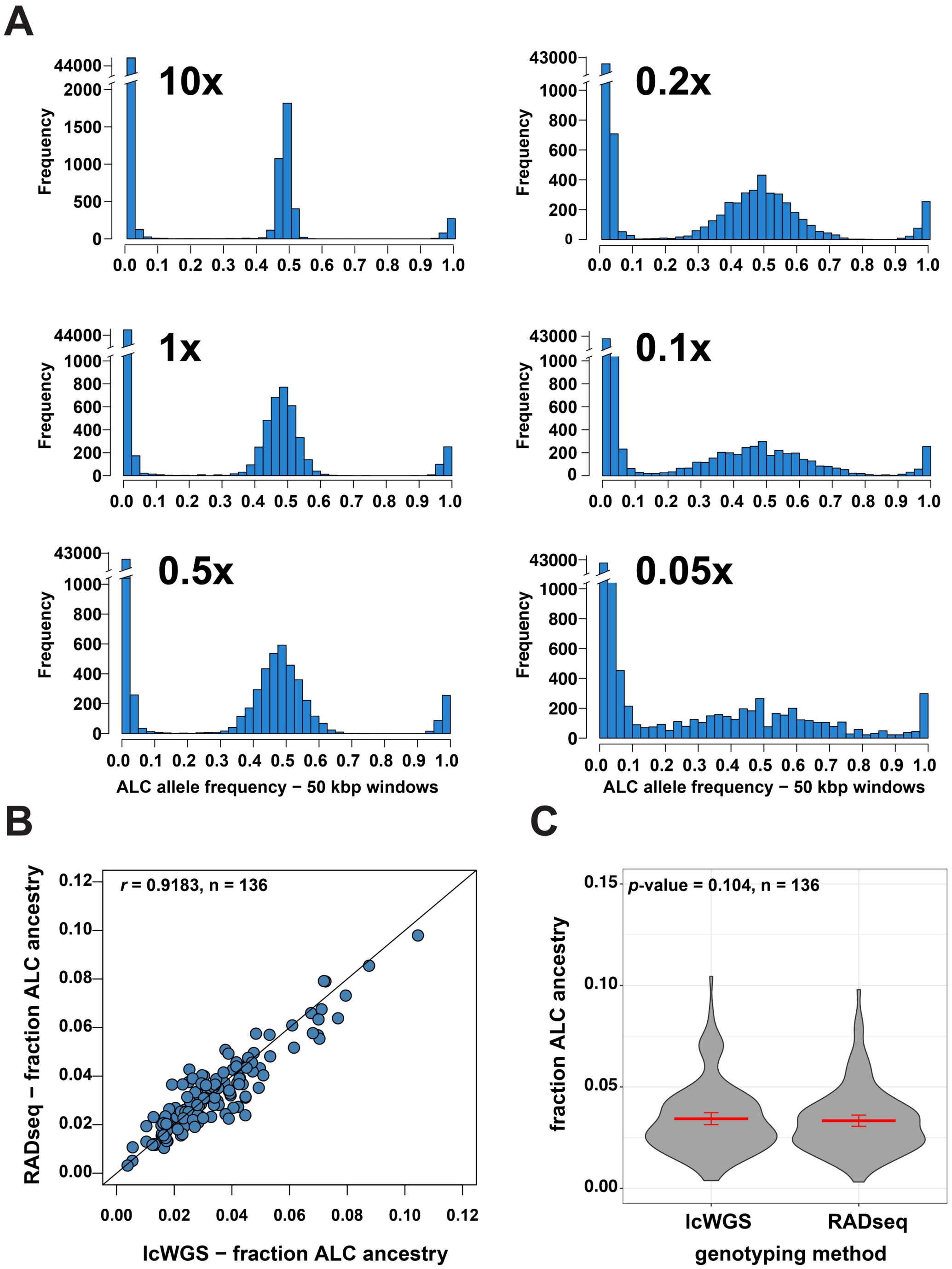


**Figure S1 | Leopard cat ancestry inference from WGS – (A)** Distributions of leopard cat allele count fractions, summed for ancestry informative loci in 48,208 50kb windows across the domestic cat assembly (felCat9). WGS from a representative Bengal cat, with coverage depth downsampled to determine minimum coverage depths for ancestry inference. **(B)** A scatter plot of genome-wide leopard cat ancestry estimated from RADseq and lcWGS data for 136 Bengal cats. **(C)** Violin plots showing the distributions, means, and standard errors (lcWGS: 3.43±0.15%, RADseq: 3.33±0.15%; Student's t-test *P* = 0.104) for genome-wide leopard cat ancestry estimated from lcWGS or RADseq data.

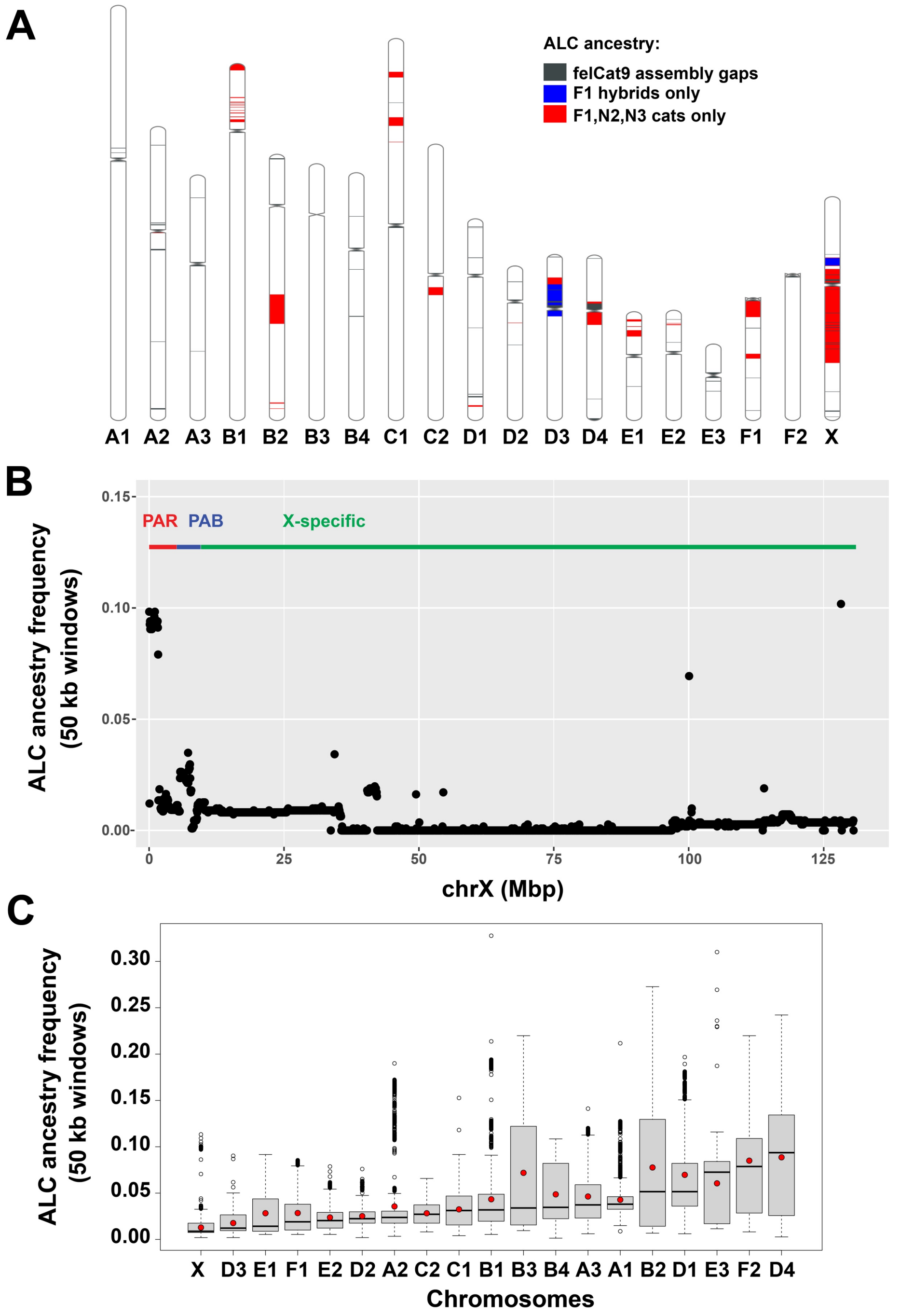


**Figure S2 | Genomic intervals without leopard cat ancestry the Bengal cat breed - (A)** A domestic cat ideogram (felCat9) depicting intervals devoid of leopard ancestry in F1 hybrid offspring (n=730, blue regions) or SBT Bengal cats (n=702, red regions). Grey regions represent intervals with assembly gaps where ancestry inference was not possible. **(B)** Leopard cat haplotype frequencies across the X chromosome, including the pseudoautosomal region (PAR), the pseudoautosomal boundary (PAB), and the X specific segments. **(C)** Boxplots of leopard cat ancestry fraction from 702 SBT Bengal cats, measured in 50kb windows across each chromosome. The median (black line) and mean (red circle) leopard cat ancestry fraction for all 50kb windows are indicated.

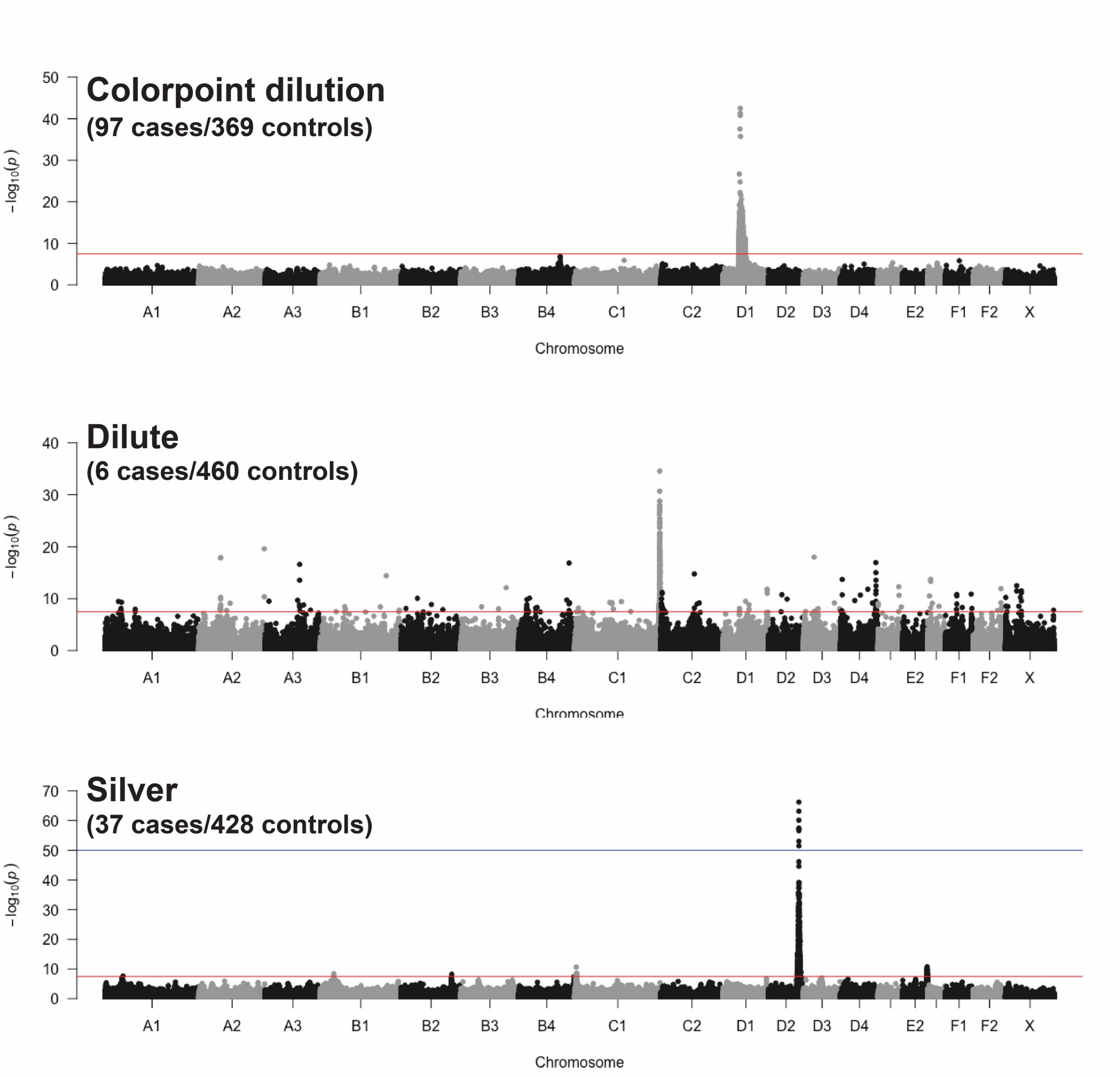


**Figure S3 | Mapping coat color traits in the Bengal cat breed with impute lcWGS -** Manhattan plots of GWAS with 1,583,065 SNVs imputed from lcWGS for three Mendelian coat color traits segregating in the Bengal breed.

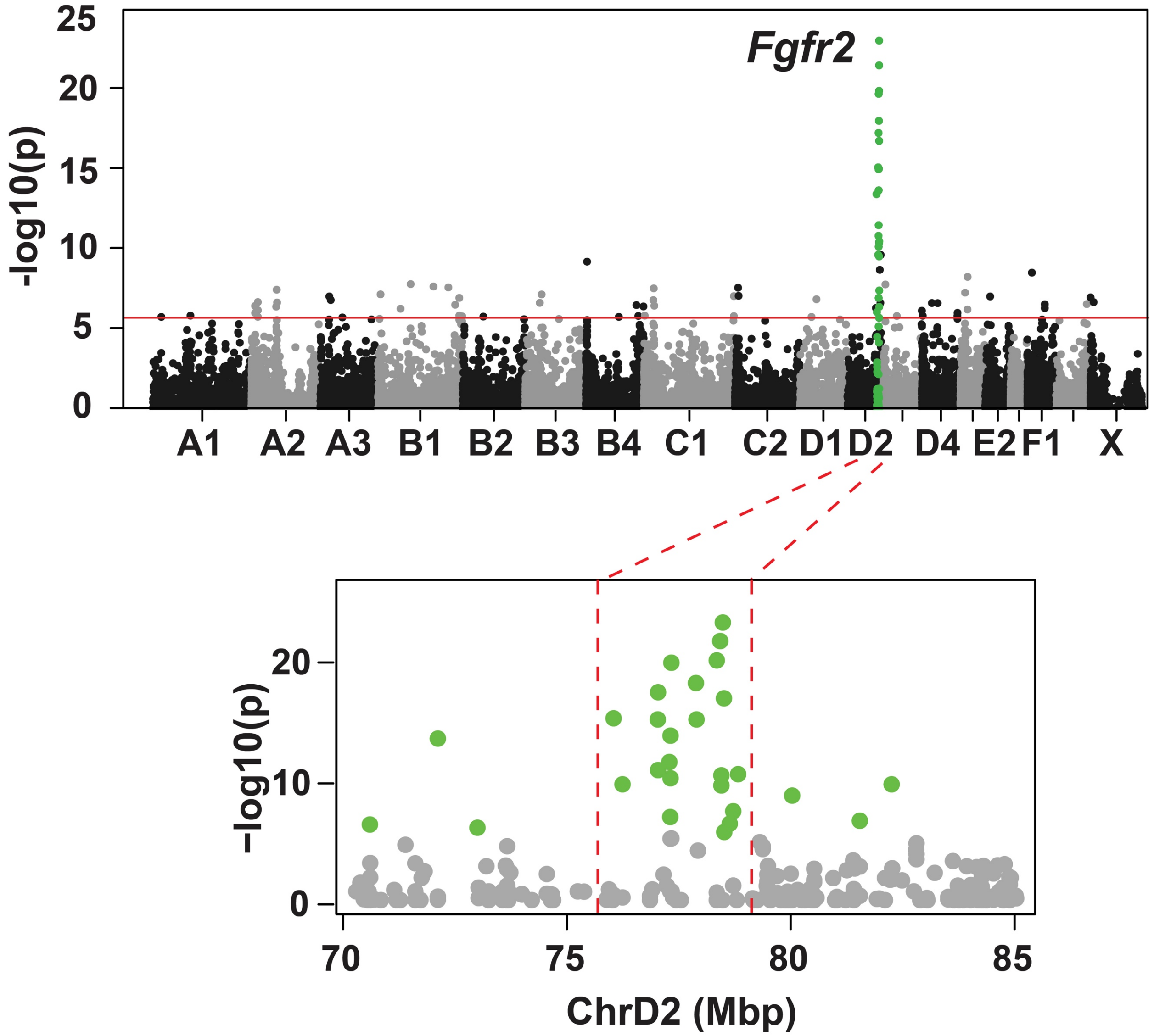


**Figure S4 | Glitter GWAS from RADseq-genotyped Bengal cats -** Top panel - a case-control GWAS to identify a *Glitter* association interval (green), using 66,289 RADseq-derived SNVs, with an association peak at chrD2:78,392,945 (Fischer exact test $P = 9.0 \times 10^{-23}$). Bottom panel – a higher resolution view of the *Glitter* association. Broken red lines delineate a 2.47 Mb *Glitter* association interval (48 RADseq SNVs), depicted in Figure 6D.

**Table S1: Intronic *Fgfr2* LTR insertion genotypes in glitter and non-glitter Bengal cats**

| Glitter status | *LTR/LTR* | *LTR/+* | +/+ |
|---|---|---|---|
| glitter | 80 | 0 | 0 |
| non-glitter | 0 | 33 | 7 |
